# Trace-DNA from a century-old holotype specimen resolves taxonomic uncertainties: the case of the Hawaiian pink precious coral (*Pleurocorallium secundum*), a CITES-listed species used in jewelry

**DOI:** 10.1101/2025.03.06.641805

**Authors:** Bertalan Lendvay, Nadja V. Morf, Laurent E. Cartier, Michael S. Krzemnicki, Masanori Nonaka

**Affiliations:** Zurich Institute of Forensic Medicine, University of Zurich, Zurich, Switzerland; Swiss Gemmological Institute SSEF, Basel, Switzerland; Okinawa Churashima Research Institute, Motobu-cho, Okinawa, Japan

**Keywords:** Species identification, DNA barcodes, coral taxonomy, Coralliidae, octocoral

## Abstract

The holotype specimen holds the most authentic characteristics of a species; its features will serve as a foundation for the identification of individuals belonging to this species. The precious coral *Pleurocorallium secundum* was described in 1846 based on a colony from the Hawaiian Islands. This specimen has been preserved; however, it is decorticated and contains exclusively the axial skeleton, which has hindered its use for accurate species identification. Therefore, the species was redescribed in 1956 based on a specimen lot collected in 1902. *Pleurocorallium secundum* was considered the most frequently fished precious coral species in the second half of the 20^th^ century with landings on the scale of thousands of tons, which was followed by its listing on CITES Appendix III. Recently, the conspecificity of the holotype and the redescribed colonies was questioned and specimens labeled in the scientific literature as *P. secundum* were discovered to be phylogenetically distant from each other. To clarify the identity of *P. secundum*, we took minimally destructive samples from the century-old holotype and the redescribed colonies and applied techniques conforming to low copy number DNA analyses. DNA sequences of three mitochondrial regions were evaluated in a phylogenetic framework together with DNA sequences retrieved from freshly collected putative *P. secundum* specimens and sequences from the scientific literature. The results of this study clearly indicate that the holotype and the redescribed colonies of *P. secundum* represent the same species. Based on the specimens confirmed to be *P. secundum* with genetic evidence, the distribution area of *P. secundum* stretches from the Hawaiian Islands to the South China Sea. At the same time, our analysis uncovered both published and fresh specimens that are in fact not *P. secundum*. The latter includes the fished Makapuu coral bed in Hawaii, which used to be a significant coral fishing area. Based on the microscopic analysis of the redescribed colonies, we complement the diagnosis of *P. secundum*. This, together with our genetic results will aid the identification of coral objects present in the international jewelry trade by providing authentic molecular barcoding markers and morphologic features for the identification of *P. secundum*.

**Statements and Declarations:** The authors have no competing interests to declare that are relevant to the content of this article.

## Introduction

Within the vastly diverse class of Octocorallia, precious corals (genera *Corallium*, *Hemicorallium* and *Pleurocorallium*, Coralliidae) are a distinctive group characterized by possessing an unjointed axial medulla of solid calcium carbonate with bright colors (Bayer 1956). This hard and colorful internal skeletal axis has made precious corals a highly valuable material for the jewelry industry for centuries (Cartier *et al*. 2018). Besides the long-known precious coral fishing grounds in the Mediterranean sea and the waters off the islands of Japan and Taiwan, precious corals have been discovered in the waters surrounding the Hawaiian Islands and the Emperor Seamounts in 1965 (Grigg 2002; Tsounis *et al*. 2010). This region has become the most important coral fishing area for decades (Bruckner 2009). The pink and white-pink spotted corals of the North Pacific originate from several *Hemicorallium* and *Pleurocorallium* species, among them the one harvested in the largest quantities, the Hawaiian pink coral, *Pleurocorallium secundum* (Grigg 1993; Parrish *et al*. 2009; Parrish *et al*. 2017; Nonaka & Hayashibara 2021). In the trade and jewelry industry, pink colored Hawaiian corals are generally associated with this species and are known by several trade names, in particular Midway, pink, white/pink (bianco rosa), angel skin, boké, middo sango and rosato coral (Tsounis *et al*. 2010; Cooper *et al*. 2011; CIBJO 2017, 2020).

*Pleurocorallium secundum* was described by James D. Dana from the Hawaiian Islands with the former name *Corallium secundum* in 1846 as the second precious coral species in the Coralliidae family following the type species, *Corallium rubrum* (as *Madrepora rubra*, Linnaeus 1758) (Dana 1846). It was first mentioned as *P. secundum* by Gray (1867), and its assignment to the *Pleurocorallium* genus has been later supported by phylogenetic studies (Ardila *et al*. 2012; Tu *et al*. 2015). The *P. secundum* specimen illustrated in Dana’s species description has been preserved in the Smithsonian National Museum of Natural History, Washington D.C. (USNM) with the lot number USNM 600 and is recognized since as the holotype specimen (Figure 1A, B; Bayer 1956; Tu *et al*. 2016). Dana described characteristics of the cortex, however already on Dana’s drawing, the specimen seems to completely lack the cortex as if it was meticulously removed and merely possess the axial medulla (Figure 1C). For over a century, this specimen was the single colony known from the species. Then, in 1956 Fredrick Bayer identified *P. secundum* colonies from the material of the “Albatross” research vessel’s Hawaiian expedition in 1902 (Bayer 1956). Moreover, he redescribed the species to amend its identification key to conform to other, later described morphologically similar species. Due to incompleteness of the holotype specimen, Bayer wrote the new diagnosis based on two colonies of the “Albatross” material. The data record for one of these colonies matches with those of lot USNM 49930 (Figure 1D), which was considered *P. secundum*’s “neotype” by Nonaka *et al*. (2014). However, as Bayer did not designate these colonies as a neotype, in the following, we refer to it as “redescribed colonies”. Nonaka *et al*. (2014) carefully studied Dana’s holotype (USNM 600) and Bayer’s redescribed colonies (USNM 49930). They concluded that due to Dana’s vague species description and the incompleteness of the holotype specimen, there is “no objective evidence that USNM 600 and USNM 49330 may be or in fact are the same species, *C. secundum*”. This implies that “if Bayer’s specimen (USNM 49330) is a different species than Dana’s specimen (USNM 600), then USNM 49330 needs to be described as a new species. In this case, almost all specimens heretofore identified as *C. secundum* would have to be re-assigned to this new species.”

**Fig. 1.**
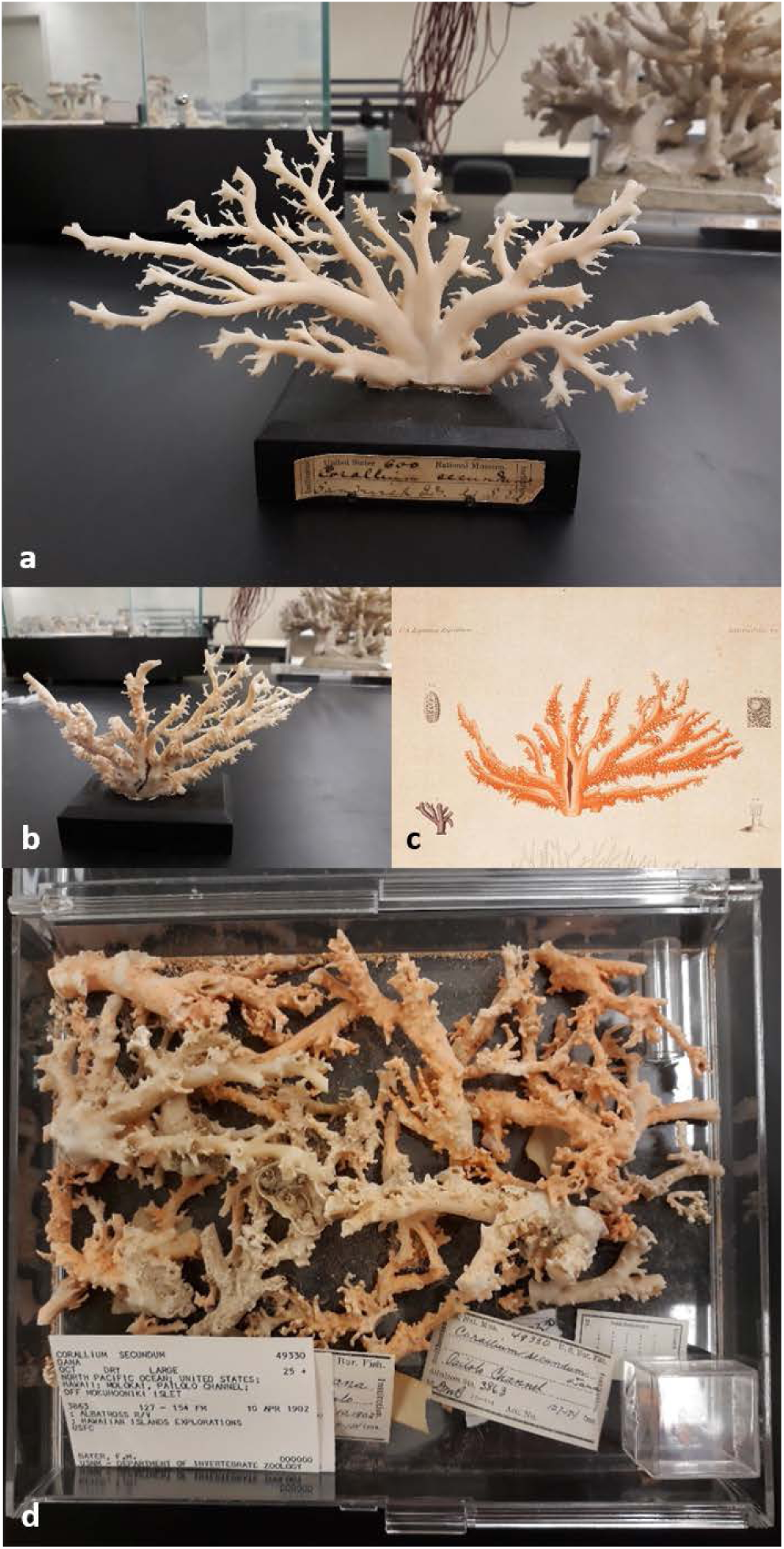
The holotype and the redescribed colonies of *Pleurocorallium secundum* at the dry coral collection of the Smithsonian National Museum of Natural History. The holotype specimen is photographed from the labeled “front” side (a) and from an angle (b) similar from which Dana depicted the specimen in his original species description from 1846 (c). The lot of the redescribed colonies (d) contains multiple fragments collected in 1902

Besides the upper outlined taxonomic uncertainties, there is an additional identification problem with specimens of *P. secundum*. Two phylogenetic studies, Ardila *et al*. (2012) and Tu *et al*. (2015) analyzed altogether 14 *P. secundum* specimens of the USNM collection (Table 1). In both studies, these 14 specimens clustered in two clearly distinct genetic groups: 11 specimens clustered together with other Hawaiian specimens labeled as *Pleurocorallium niveum*. The remaining three specimens formed a well-supported genetic cluster and were considered *P. secundum*. These few samples served as reference samples for Lendvay *et al*. to characterize *P. secundum* for their genetic species identification assay called *Coral*-ID (2022). It was however noticed that based on this method, the single *P. secundum* complete mitochondrial genome on NCBI GenBank (accession number KC782347) from the study of Figueroa and Baco (2014) would classify as another *Pleurocorallium* species (B. Lendvay, unpublished data).

**Table 1.**
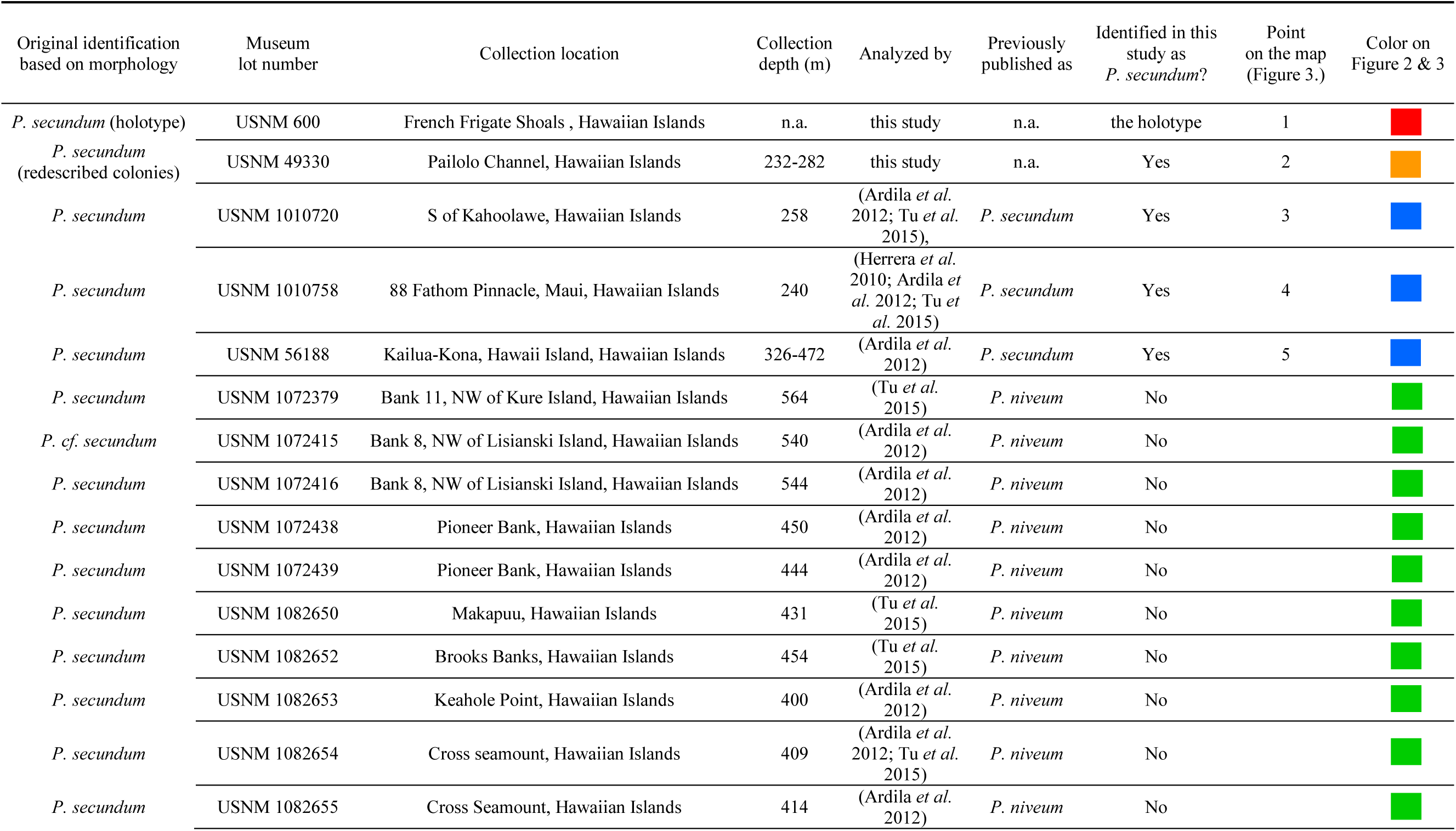
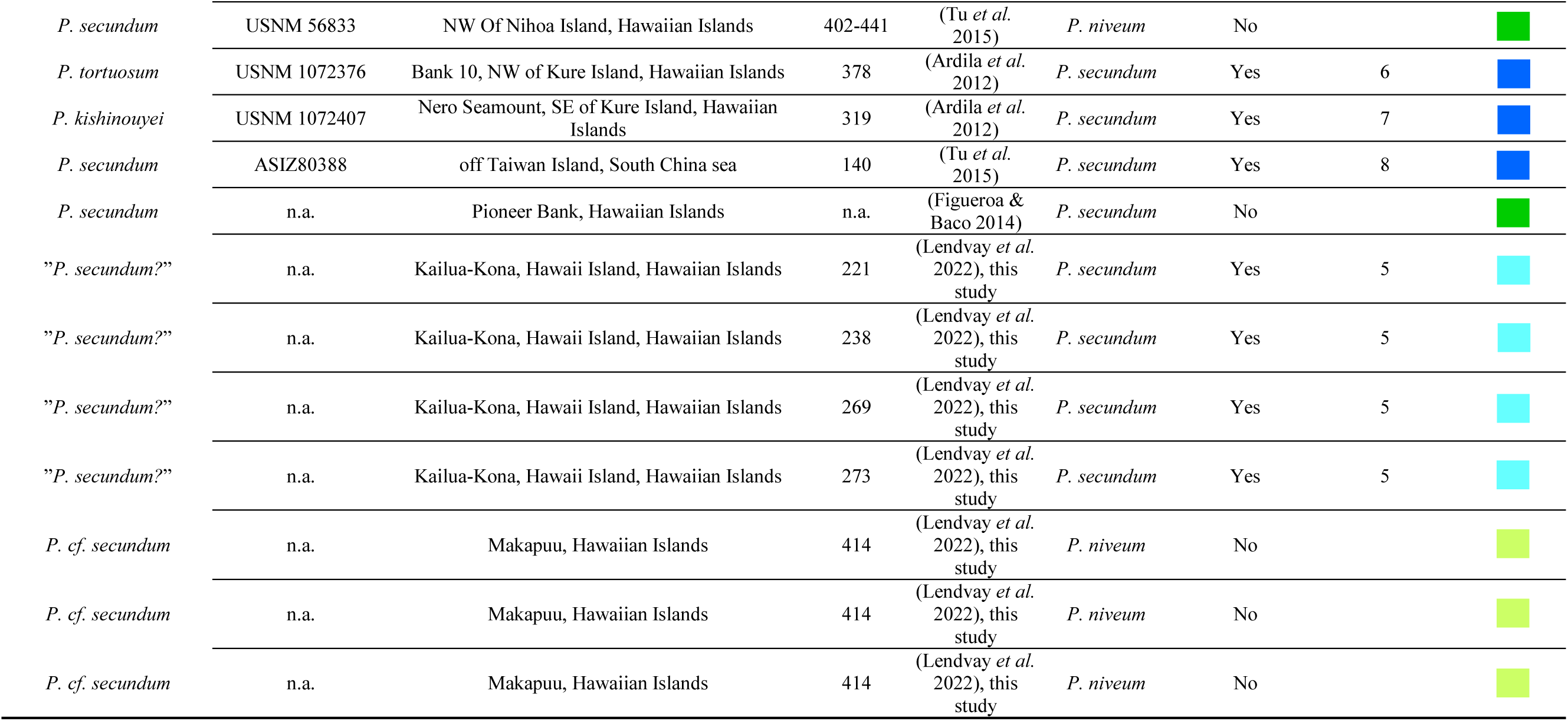
All specimens that have been considered *Pleurocorallium secundum* before and/or after genetic analysis to this date. Colored squares represent the following: red: holotype; orange: the redescribed colonies; light blue and dark blue: respectively fresh samples and samples from the literature identified as *P. secundum*; light green and dark green: respectively fresh samples and samples from the literature originally identified as *P. secundum* but disproven here. Note that the *P. niveum* specimens marked with dark green squares were originally identified as *P. secundum* but already considered *P. niveum* by Tu *et al*. (2015), which is confirmed here.

| Original identification based on morphology | Museum lot number | Collection location | Collection depth (m) | Analyzed by | Previously published as | Identified in this study as *P. secundum*? | Point on the map (Figure 3.) | Color on Figure 2 & 3 |
| --- | --- | --- | --- | --- | --- | --- | --- | --- |
| *P. secundum* (holotype) | USNM 600 | French Frigate Shoals , Hawaiian Islands | n.a. | this study | n.a. | the holotype | 1 |  |
| *P. secundum* (redescribed colonies) | USNM 49330 | Pailolo Channel, Hawaiian Islands | 232-282 | this study | n.a. | Yes | 2 |  |
| *P. secundum* | USNM 1010720 | S of Kahoolawe, Hawaiian Islands | 258 | (Ardila *et al.* 2012; Tu *et al.* 2015), | *P. secundum* | Yes | 3 |  |
| *P. secundum* | USNM 1010758 | 88 Fathom Pinnacle, Maui, Hawaiian Islands | 240 | (Herrera *et al.* 2010; Ardila *et al.* 2012; Tu *et al.* 2015) | *P. secundum* | Yes | 4 |  |
| *P. secundum* | USNM 56188 | Kailua-Kona, Hawaii Island, Hawaiian Islands | 326-472 | (Ardila *et al.* 2012) | *P. secundum* | Yes | 5 |  |
| *P. secundum* | USNM 1072379 | Bank 11, NW of Kure Island, Hawaiian Islands | 564 | (Tu *et al.* 2015) | *P. niveum* | No |  |  |
| *P. cf. secundum* | USNM 1072415 | Bank 8, NW of Lisianski Island, Hawaiian Islands | 540 | (Ardila *et al.* 2012) | *P. niveum* | No |  |  |
| *P. secundum* | USNM 1072416 | Bank 8, NW of Lisianski Island, Hawaiian Islands | 544 | (Ardila *et al.* 2012) | *P. niveum* | No |  |  |
| *P. secundum* | USNM 1072438 | Pioneer Bank, Hawaiian Islands | 450 | (Ardila *et al.* 2012) | *P. niveum* | No |  |  |
| *P. secundum* | USNM 1072439 | Pioneer Bank, Hawaiian Islands | 444 | (Ardila *et al.* 2012) | *P. niveum* | No |  |  |
| *P. secundum* | USNM 1082650 | Makapuu, Hawaiian Islands | 431 | (Tu *et al.* 2015) | *P. niveum* | No |  |  |
| *P. secundum* | USNM 1082652 | Brooks Banks, Hawaiian Islands | 454 | (Tu *et al.* 2015) | *P. niveum* | No |  |  |
| *P. secundum* | USNM 1082653 | Keahole Point, Hawaiian Islands | 400 | (Ardila *et al.* 2012) | *P. niveum* | No |  |  |
| *P. secundum* | USNM 1082654 | Cross seamount, Hawaiian Islands | 409 | (Ardila *et al.* 2012; Tu *et al.* 2015) | *P. niveum* | No |  |  |
| *P. secundum* | USNM 1082655 | Cross Seamount, Hawaiian Islands | 414 | (Ardila *et al.* 2012) | *P. niveum* | No |  |  |
| *P. secundum* | USNM 56833 | NW Of Nihoa Island, Hawaiian Islands | 402-441 | (Tu *et al.* 2015) | *P. niveum* | No |  |  |
| *P. tortuosum* | USNM 1072376 | Bank 10, NW of Kure Island, Hawaiian Islands | 378 | (Ardila *et al.* 2012) | *P. secundum* | Yes | 6 |  |
| *P. kishinouyei* | USNM 1072407 | Nero Seamount, SE of Kure Island, Hawaiian Islands | 319 | (Ardila *et al.* 2012) | *P. secundum* | Yes | 7 |  |
| *P. secundum* | ASIZ80388 | off Taiwan Island, South China sea | 140 | (Tu *et al.* 2015) | *P. secundum* | Yes | 8 |  |
| *P. secundum* | n.a. | Pioneer Bank, Hawaiian Islands | n.a. | (Figueroa & Baco 2014) | *P. secundum* | No |  |  |
| "*P. secundum*?" | n.a. | Kailua-Kona, Hawaii Island, Hawaiian Islands | 221 | (Lendvay *et al.* 2022), this study | *P. secundum* | Yes | 5 |  |
| "*P. secundum*?" | n.a. | Kailua-Kona, Hawaii Island, Hawaiian Islands | 238 | (Lendvay *et al.* 2022), this study | *P. secundum* | Yes | 5 |  |
| "*P. secundum*?" | n.a. | Kailua-Kona, Hawaii Island, Hawaiian Islands | 269 | (Lendvay *et al.* 2022), this study | *P. secundum* | Yes | 5 |  |
| "*P. secundum*?" | n.a. | Kailua-Kona, Hawaii Island, Hawaiian Islands | 273 | (Lendvay *et al.* 2022), this study | *P. secundum* | Yes | 5 |  |
| *P. cf. secundum* | n.a. | Makapuu, Hawaiian Islands | 414 | (Lendvay *et al.* 2022), this study | *P. niveum* | No |  |  |
| *P. cf. secundum* | n.a. | Makapuu, Hawaiian Islands | 414 | (Lendvay *et al.* 2022), this study | *P. niveum* | No |  |  |
| *P. cf. secundum* | n.a. | Makapuu, Hawaiian Islands | 414 | (Lendvay *et al.* 2022), this study | *P. niveum* | No |  |  |

*Pleurocorallium secundum* was fished between the 1960s and 1990s in quantities estimated to reach the scale of hundred tons (Cannas *et al*. 2019). Subsequently, its harvesting around the Midway atoll and the Hawaiian Archipelago decreased dramatically due to the declining price of corals and resource depletion and its commercial harvest ceased completely in 2001 (Grigg 2002; Chang *et al*. 2013). However, the harvesting of *P. secundum* reportedly continued in the South China sea and Philippine sea in small quantities, and *P. secundum* has been placed on the CITES Appendix III together with three other precious coral species by China as of July 2008 (Shiraishi 2018). A criterion of including a species in CITES Appendix III is that the species is native in the submitting country. Due to the lack of a sound taxonomic identification of Chinese corals (fished off Taiwan) as *P. secundum*, the report published by the United Nations FAO in 2019 raised concerns over the legitimacy of the inclusion of *P. secundum* in CITES Appendix III by China (Cannas *et al*. 2019).

Given all the uncertainties surrounding the identity of *P. secundum*, we considered it timely to re-evaluate its taxonomic status. Based on the genetic analysis of the holotype and redescribed specimens, fresh samples and published data from the literature, we aimed to answer the following specific questions: i) Do the holotype and the redescribed colonies of *P. secundum* in fact belong to the same species? ii) Is it possible to establish solid species-characteristic morphologic attributes and DNA barcodes for the identification of *P. secundum*? iii) What is the geographical distribution of *P. secundum* and can we confirm that the corals considered *P. secundum* fished in Chinese waters really belong to this species?

## Methods

### Analyzed coral samples

We analyzed three types of samples in a phylogenetic framework: *i)* the holotype and the redescribed colonies of *P. secundum*, *ii)* fresh samples of putatively *P. secundum* samples collected in two areas off the Southeastern Hawaiian Islands and *iii)* DNA sequence data of samples from the scientific literature - including data from *P. secundum* from Chinese waters collected by the Taiwanese precious coral fishery.

Holotype and the redescribed colonies: The holotype and the redescribed colonies of *P. secundum* are held in the dry coral collection of the Smithsonian National Museum of Natural History, Washington D.C. and were sampled in November 2022. The holotype specimen, lot number USNM 660, is approximately 20 cm wide and 10 cm high and glued on a wooden base. Two samples were collected from this specimen for genetic analyses; the first sample, approximately 3 mg in weight, was skeletal material scraped off the surface at the base of the specimen with a scalpel. The second sample contained formerly broken off skeletal buds with a total weight of approximately 5 mg. The lot of the redescribed colonies, lot number USNM 49330, contains ca. 25 branches, the largest ones approximately 7-8 cm long. The lot box also contains considerable amount of fallen off coenenchymal debris. About 20 mg of bulk debris was collected for genetic testing. Bayer (Cannas *et al*. 2019) described the morphology of these specimens in detail, but did not mention the size of the sclerites. Therefore, small amounts of coenenchyme were taken from four colony parts (tentacles, autozooid mounds, branch tips, and colony base) on the specimen for scanning electron microscope (SEM) imaging of the sclerites. The sclerites were separated and cleaned using 5% sodium hypochlorite solution (household bleach), and details of them were observed with a Keyence VE-8800 instrument. Their length and widths were measured with SEM accessory software.

Fresh samples: We analyzed seven fresh samples presumed to be *P. secundum* from two locations of the Hawaiian Islands. Three samples were collected in July 2013 at 414 m depth at the Makapuu coral bed located between Oahu and Molokai islands. These samples were later considered *P. niveum* by Lendvay *et al*. (2022). Four additional samples originated from the coast of Kailua-Kona, Hawaii Island and were collected in September 2016 at the depths between 221 and 273 m. These samples were considered *P. secundum* by Lendvay *et al*. (2022) and were named “*P. secundum*?” in the publication by Conner *et al*. (2023). The fresh samples were collected using Pisces IV/V submersibles during Hawaii Undersea Research Laboratory (HURL) cruises, and stored and transported in ethanol solution.

Data from scientific literature: We searched all *Pleurocorallium* specimens that had DNA sequences available from three variable mitochondrial DNA regions; the mtND6-COI intergenic spacer (IGS), a fragment of the DNA mismatch repair protein (MutS) gene and a fragment of the large ribosomal RNA gene subunit (LR) on the NCBI GenBank platform. On 1^th^ November 2023, a total of 52 specimens were found and sequence data of the three genes were downloaded (see the detailed data set in Supporting Information S2). These included 48 specimens analyzed by Tu *et al*. (2015), two specimens from Figueroa and Baco (2014) and Uda *et al*. (2013), respectively, and one specimen analyzed by Uda *et al*. (2011). These samples included four specimens labeled as *P. secundum*. The scientific literature was then reviewed in order to discover any other scientific works that analyzed these four specimens (based on DNA sequences of other genes).

### Laboratory protocols

DNA was extracted from the skeletal material of the holotype specimen by applying a protocol specially optimized for DNA extraction from minute amounts of coral axis powder. The method is based on the complete decalcification and lysis of the skeletal powder followed by a column purification of the lysate and has previously proved to yield DNA from as low as 2.3 mg material drilled from polished coral twigs (Lendvay *et al*. 2020; Lendvay *et al*. 2022). The coenenchymal debris of the redescribed colonies and the coenenchymal tissue of the fresh samples were extracted using the QiaAmp DNA Mini kit (Qiagen) according to the manufacturer’s protocol.

For all samples, we analyzed the DNA sequences of the three mitochondrial DNA regions, IGS, MutS and LR. To amplify the fragmented DNA of the holotype and redescribed colonies, for both IGS and MutS, we designed primers to amplify three overlapping short fragments of any *Pleurocorallium* species. The sequences of the newly designed IGS and MutS primers, PCR protocols, amplicon lengths and the positions of the amplified IGS and MutS fragments (as mapped on the *Pleurocorallium konojoi* mitochondrial reference genome (NC_015406, Uda *et al*. 2011) are shown in Supporting information S1. The amplified short IGS and MutS fragments were sequenced by Sanger technology according to Lendvay *et al*. (2022). For the fresh samples that have less fragmented DNA, the IGS and MutS regions were amplified and sequenced as single fragments following Tu *et al*. (2015). The very short LR region was amplified for all samples using the same primers previously applied by Lendvay *et al*. (2020), see Supporting information S1. The LR amplicons were then converted to sequencing libraries using the KAPA HyperPrep PCR-free Kit (Roche) and run on a MiSeq instrument (Illumina).

DNA extracted from the holotype and redescribed colonies were PCR-amplified and sequenced twice to produce independent technical replicates in ISO17025-accredited trace-DNA laboratory rooms of the Institute of Forensic Medicine at the University of Zurich. The DNA of the fresh samples was amplified and sequenced in separate research-purpose laboratory rooms at the same facility.

### DNA sequence analysis

For the IGS and MutS datasets, primer sequences were trimmed from the raw DNA sequence data. Then, in the case of the holotype and the redescribed colonies that were amplified as short overlapping DNA fragments, the short fragments were assembled to yield the complete analyzed IGS and MutS fragments. The LR sequence data was evaluated as described by Lendvay *et al*. (2020).

The DNA sequences obtained by the two technical replicates (for all three analyzed DNA regions: IGS, MutS and LR) performed respectively for the DNA extracts of both holotype and redescribed colonies were aligned to each other to test for consistency of the replicate data. In the following, the DNA sequence data of the *P. secundum* holotype, the redescribed colonies, the fresh Hawaiian coral samples and the *Pleurocorallium* sequence data from the literature were aligned and trimmed together with the IGS, MutS and LR sequences of the *Corallium rubrum* mitochondrial reference genome (NCBI GenBank accession number AB700136, Uda *et al*. 2013), which was used as outgroup. Trimming, assembly and alignment of the DNA sequences were done in Geneious Prime 2022.0.2 (www.geneious.com). Bayesian phylogenetic trees were created with partitioning the concatenated alignment of the three genes and run in MrBayes version 3.2.7 (Ronquist *et al*. 2012) with the settings used by Lendvay *et al*. (2020): mixed substitution model prior, 10^7^ generations with sampling every 10^3^ generations and 25 % burn-in. A majority rule consensus tree was constructed from the raw tree data in FigTree (Rambaut 2012). Based on their DNA sequence similarity and phylogenetic position compared to the holotype specimen, we re-evaluated whether specimens formerly identified as *P. secundum* indeed belong to this species. We then pinned the location of the confirmed specimens on a map to estimate the approximate geographical distribution of *P. secundum*.

## Results

The SEM images and sizes of the redescribed colonies’ (USNM49930) sclerites are shown on Fig. 4 and Table 2, respectively. The type of sclerites fit Bayer’s description. Tentacles contained many 8-radiates, while spiny rods were absent. In the coenenchyme, double-clubs were dominant, and 6-, 7-, 8-radiates were not common. However, in this study, unlike in Bayer’s description, many asymmetric 8-radiates appeared.

**Table 2.**
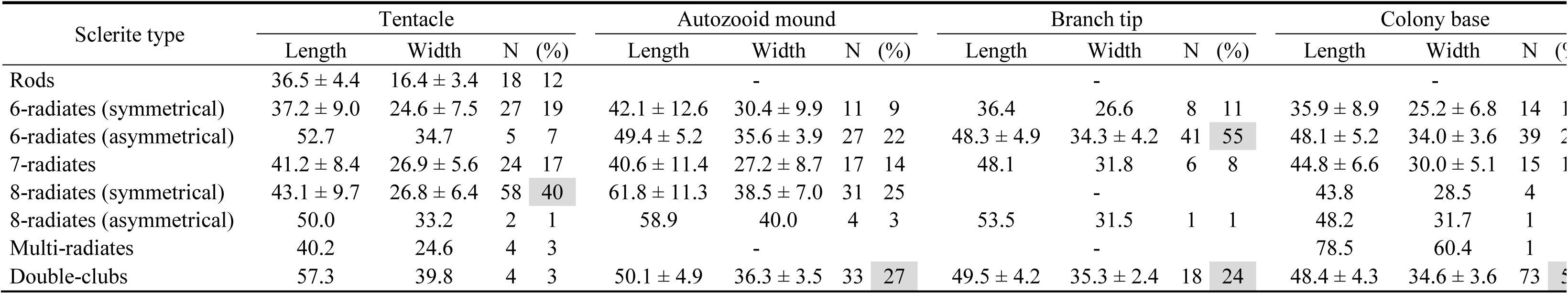
Summary of length and width measurements of sclerites of *Pleurocorallium secundum* USNM49330. Measurements are reported as the average ± standard deviation (more than 10%), or only the average (less than 9%), in mm. Numbers with gray background indicate most abundant sclerite type. “-” means not found.

Two technical replicates were performed for each of the seven DNA fragments for both the holotype and the redescribed colonies. In all cases, these replicates resulted in identical DNA sequences, which testifies the accuracy of the results. The IGS, MutS and LR DNA sequences were respectively 487, 400 and 127 bp long in all newly sequenced samples. The NCBI GenBank accession numbers and the corresponding aligned sequences are presented in Supporting Information S2.

The IGS and LR sequences were identical in the holotype and the redescribed colonies. The MutS region contained a single difference (T in the holotype, C in the redescribed colonies at the position of base 7221 in the *P. konojoi* mitochondrial reference genome); this third-codon base difference results in a same-sense mutation. The combined IGS-MutS-LR alignment was 1090 bp in total when including *C. rubrum* as outgroup, which introduced numerous indels in the alignment (see alignments in Supporting Information S2).

The *P. secundum* holotype and the redescribed colonies created a highly supported phylogenetic branch (with posterior probability value of 1) together with the four fresh specimens *P. secundum* from Kailua-Kona, Hawaii Island and the three *P. secundum* specimens from the study of Tu *et al*. (2015; specimens marked HW_1, HW_2 and TW on Figure 2). All samples clustering with the holotype specimen were identical with the redescribed colonies, sharing that single base difference compared to the holotype (99.9% sequence similarity compared to the *P. secundum* holotype). Any other analyzed *Pleurocorallium* sample comprised less than 97.3% sequence similarity compared to the *P. secundum* holotype. The fresh specimens from Makapuu as well as the *P. secundum* sample sequenced by Figueroa and Baco (2014) grouped together with specimens formerly found to be *P. niveum* from the Hawaiian Islands and *P. bonsaiarborum* from New Caledonia. We assessed whether the specimens previously considered *P. secundum* were correctly identified based on their phylogenetic position and genetic similarity to the *P. secundum* holotype (Table 1). We considered all specimens clustering together with the *P. secundum* holotype (with 99.9% sequence similarity) to be indeed *P. secundum* (marked orange, dark blue and light blue on Table 1 and Figure 2). The collection location of these specimens were marked on a map using the same color code (Figure 3). All other specimens were considered to originate from a different species.

**Fig. 2.**
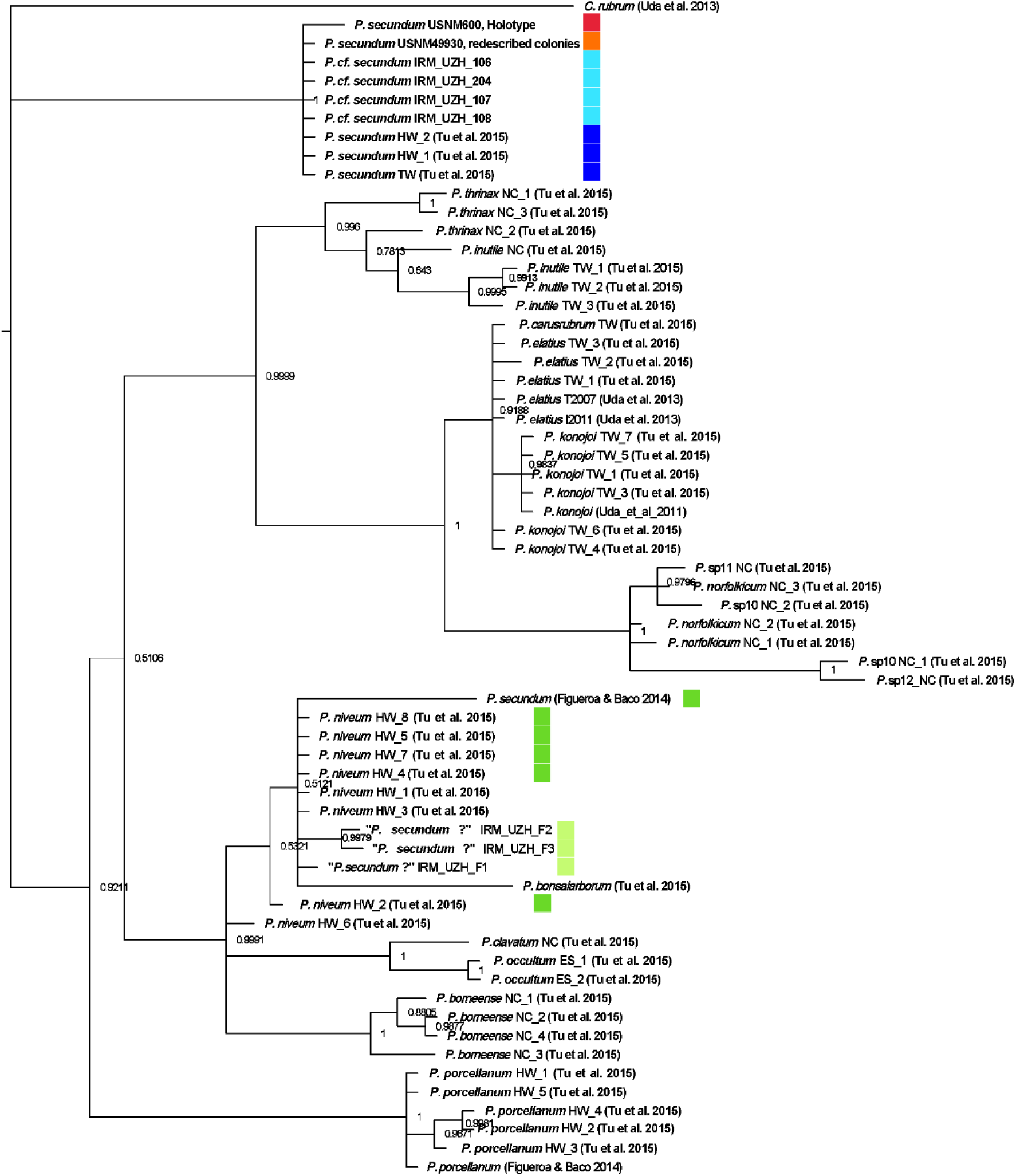
Bayesian phylogenetic tree of the holotype and the redescribed colonies of *P. secundum* together with fresh samples and published *Pleurocorallium* data based on the concatenated dataset of three mitochondrial regions. Scientific names are followed by colony identifiers and literature references where applicable. Posterior probability values are displayed on each node. Colored squares represent the following: red: holotype; orange: the redescribed colonies; light blue and dark blue: respectively fresh samples and samples from the literature identified as *P. secundum*; light green and dark green: respectively fresh samples and samples from the literature originally identified as *P. secundum* but disproven here. Note that the *P. niveum* specimens marked with dark green squares were originally identified as *P. secundum* but already considered *P. niveum* by Tu *et al*. (2015), which is confirmed here

**Fig. 3.**
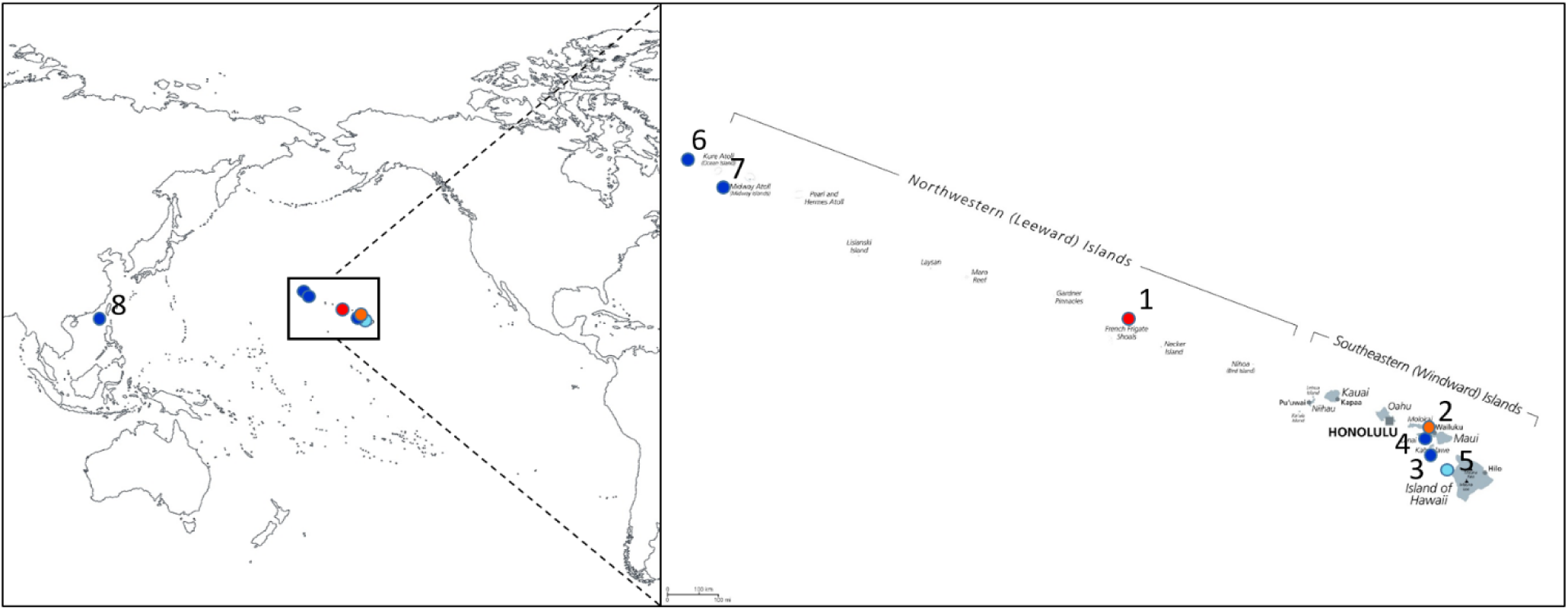
The geographic distribution of *Pleurocorallium secundum* based on specimens confirmed by genetic analyses. Colors represent the following: red: holotype; orange: the redescribed colonies; light blue: fresh samples; dark blue: museum specimens

**Fig. 4.**
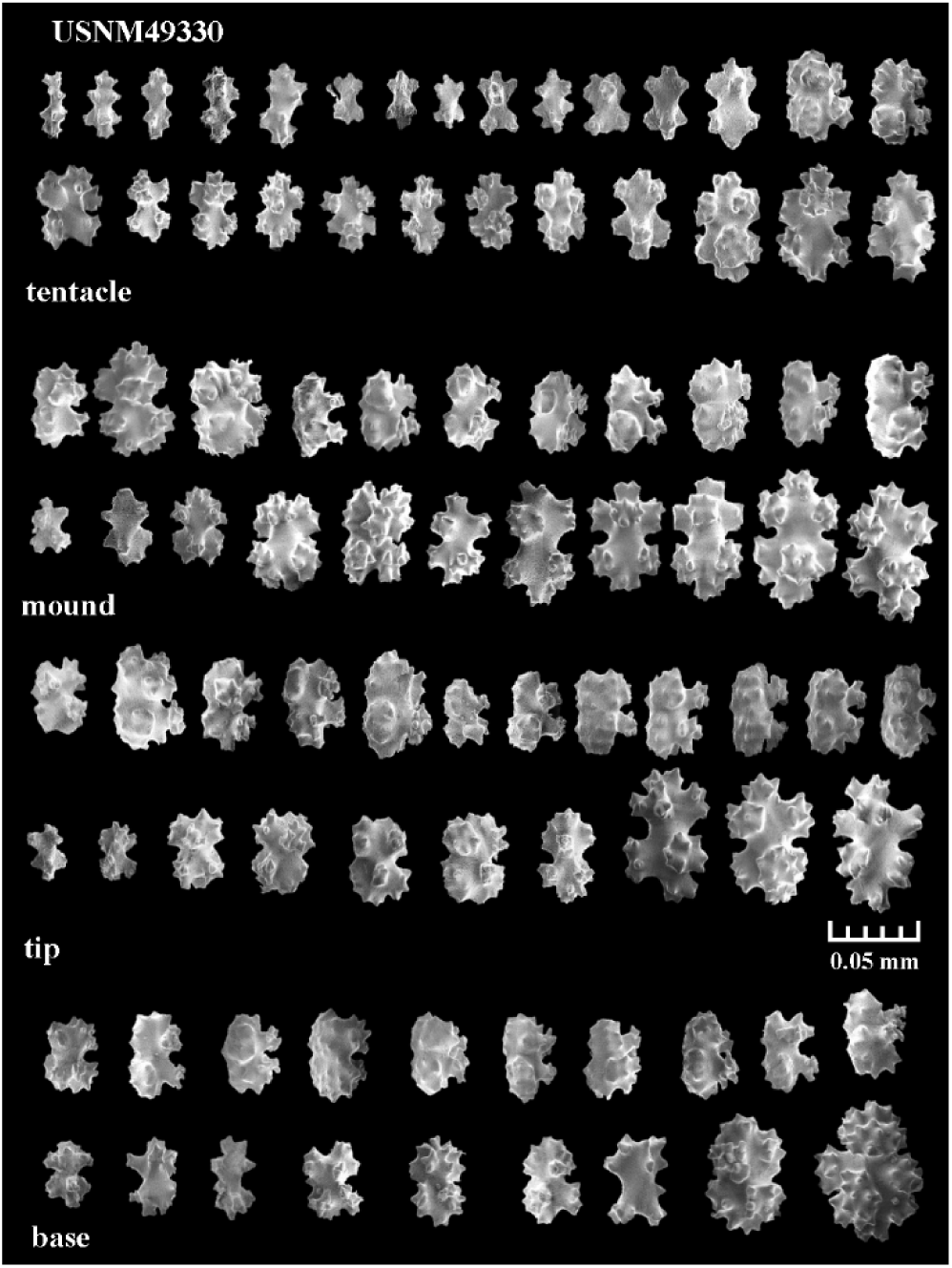
Scanning electron microscopy images of the sclerites of the redescribed colonies of *Pleurocorallium secundum* (USNM49330)

## Discussion

The genetic analysis of museum specimens has become routine in taxonomy and evolutionary biology (Raxworthy & Smith 2021). Type specimens would constitute the best reference material for species identification and the scientific value of DNA reference databases would be greatly enhanced if species were also represented by sequences of the respective type material, especially the holotype (Schäffer *et al*. 2017). However the usage of holotypes in molecular systematics and taxonomy is scarce and frequently not even attempted, mostly due to their antiquity and preservation history (Erpenbeck *et al*. 2016). Nevertheless, holotype specimens have been used for elucidating taxonomic relations of various animal taxa e.g. sponges (Erpenbeck *et al*. 2016), fishes (Agne *et al*. 2022; Sullivan *et al*. 2022), amphibians (Rancilhac *et al*. 2020; Goutte *et al*. 2022), birds (Kirchman *et al*. 2010) or mammals (Castañeda-Rico *et al*. 2022; Nations *et al*. 2022). On a single occasion, a precious coral species’ type specimen was also genotyped (*Corallium medea* from the Bahamas, Ardila *et al*. 2012).

The genetic study of a holotype is especially beneficial if the morphological characteristics allowing sound taxonomic identification are lost, which is the case with the *P. secundum* holotype. At least since 1846, this specimen exists without its cortex – possessing the most important morphological features. This also means that the material generally used for genetic analysis of precious corals (polyps or less frequently cortex e.g. France *et al*. 1996; Bayer & Cairns 2003; Aurelle *et al*. 2011; Ledoux *et al*. 2013) is lost. What remains is the skeletal axis, which is produced by a biomineralization process through the secretion of calcite crystals from an epithelial cell layer (Grillo *et al*. 1993; Perrin *et al*. 2015). Analogously to mollusk shells, the coral skeletal axis does not contain living cells and any trace-DNA molecules present in the skeletal axis are “trapped or absorbed” during the formation of the calcareous structure (Martin *et al*. 2021). Recent studies have however demonstrated that by using dedicated laboratory protocols it is feasible to perform genetic analysis even from as little as a few milligrams of precious coral axial medulla (Lendvay *et al*. 2020; Lendvay *et al*. 2022). The technical advancements in genetic analysis of minute amounts of coral skeletal material encouraged us to ask permission to sample the holotype of *P. secundum* accompanied by its redescribed colonies in order to analyze them in a phylogenetic context for resolving the taxonomic uncertainties surrounding the identity of *P. secundum*.

The nearly identical DNA sequence data of the holotype and the redescribed colonies (0.1% dissimilarity) and their genetic gap from any other haplotypes (exceeding 2.7% dissimilarity) confirm that they indeed belong to the same species. This result indicates that the identification of coral specimens as *P. secundum* based on the identification key of Bayer (Bayer 1956) and the morphologic characters of the redescribed colonies (USNM 49930) is valid. We complement Bayer’s diagnosis of the redescribed colonies by providing quantitative data and SEM images of its sclerites in the hope that these will aid accurate morphological identification of *P. secundum* colonies.

The results of this study also provide a backbone for the genetic identification of *P. secundum* by determining DNA barcode sequences from the type specimens. These barcode sequences also give us the opportunity to review specimens used in previous studies and check whether they have been correctly identified a *P. secundum*. Our results confirm that the identification of colonies as *P. secundum* by Tu *et al*. (2015) was correct. As well, the corroborating results of specimens analyzed both by Tu *et al*. (2015) and Ardila *et al*. (2012) prove that the latter authors also identified *P. secundum* specimens correctly. Likewise, the genetic classification method of Lendvay *et al*. (2022), which was based on the data of Tu *et al*. (2015) correctly identifies *P. secundum* samples and separates them from any other coral species. In turn, the specimen available to create the complete *P. secundum* mitochondrial genome (NCBI GenBank accession number KC782347) is here identified not to be *P. secundum*. As this specimen from Pioneer Bank, Hawaiian Islands clusters together with Hawaiian *P. niveum* specimens, it is likely also a member of this species.

A significant result of this study is the species confirmation of the specimen from the South China Sea as *P. secundum* (number 8 on Figure 3; P. secundum_TW, Tu *et al*. 2015). This specimen (collection accession number ASIZ80388 at the Research Museum, Biodiversity Research Center, Academia Sinica, Taipei) was collected by the Taiwanese precious coral fishery in February 2011 from the South China sea, southwest of Taiwan island. This means that the reported harvesting of *P. secundum* by Taiwanese coral fishermen (in small quantities and specifically off Lanyu Island: Shiraishi 2018; Cannas *et al*. 2019) during the 2010s is supported by genetic data from a nearby location. This result implies that the inclusion of *P. secundum* in CITES Appendix III by China was legitimate.

It is also worth to mention that the three fresh samples from Makapuu, Oahu (414 m depth) have proven not to be *P. secundum*. Besides the three specimens analyzed here, two additional specimens from Makapuu (USNM 1082650, 431 m depth, and USNM 56807, 366 m depth) have been genotyped and both turned out to be *P. niveum* (Tu *et al*. 2015). The precious coral bed off Makapuu (350-450 m depth) was discovered in 1966 and became the primary site of coral harvesting in the Southeastern Hawaiian Islands (Grigg 1993; Long & Baco 2014). Precious corals in this coral bed were previously ubiquitously considered *P. secundum* (Grigg 1988, 1993; Grigg 2002; Parrish 2007; Grigg 2010; Long & Baco 2014; Parrish *et al*. 2017). Our results raise the possibility that the fished Makapuu coral bed, at least partly, consists of a species other than *P. secundum*.

Based on the collection location of the specimens confirmed to be *P. secundum*, we can infer the approximate distribution area of the species (Figure 3.). This covers both the Hawaiian Islands and the disjunct observation in the South China Sea represented by a sole sample. The Hawaiian distribution of *P. secundum* includes the Northwestern and Southeastern Hawaiian islands from Hawaii Island to at least the 2500 km distant Kure Island. The confirmed specimens were fished at 140 m and at between 230-380 m depths in the South China Sea and the Hawaiian Islands, respectively.

## Conclusions

Identification of precious corals to the species level is often difficult even for experts. This is surely the case of coral skeletal axis fragments that are cut and polished for their use in jewelry. Lendvay *et al*. (2020) suggested minimally destructive genetic analysis as a promising tool for identification of corals from jewelry, and a protocol to distinguish CITES-listed and non-CITES-listed precious corals was developed by Lendvay *et al*. (2022). For representing *Pleurocorallium secundum*, this method relied on the MutS DNA sequence data of Tu *et al*. (2015), and the correctness of this data was confirmed by the present study. Both studies of Lendvay *et al*. (2020) and Lendvay *et al*. (2022) identified jewelry objects that were observed to originate from *Pleurocorallium* species different from those that have been known to be present in the jewelry industry (*Pleurocorallium elatius*, *P. konojoi* and *P. secundum*). The current study also confirms that these are not from *P. secundum*. To investigate the taxonomy of these *Pleurocoralliums*, reference data should be generated for the species of the genus that have no genetic data yet; *Pleurocorallium uchidai* and *P. gotoense* are both known exclusively from century-old holotypes from Japanese waters (Nonaka *et al*. 2012). A similar approach to the one used in the current study might be promising to test whether these species originating from coral fishing areas are the unknown taxa present in the coral jewelry trade.

## Supporting information

Lendvay_et_al_S1_Supporting_methods

Lendvay_et_al_S2_Supporting_results

## Acknowledgements

The authors are grateful to Andrea Quattrini, the curator of corals at the Smithsonian National Museum of Natural History for permitting the sampling of type material. We are also thankful for Samuel Kahng (University of Hawaii at Manoa) and Frank Parrish (Pacific Islands Fisheries Science Center, NOAA) for providing the fresh samples from Hawaii that were collected as part of NOAA’s Deep Coral Research and Technology Program. Frank Parrish is also acknowledged for his invaluable suggestions for improving an earlier version of the manuscript. We thank Meng-Min Hsueh (collection manager at the Biodiversity Research Museum, Academia Sinica, Taipei) for sharing information about specimens. This study was financially supported by the Emma Louise Kessler-Fonds.

## Supporting Information

**S1** Supporting Methods

A) PCR primers used for amplifying the mitochondrial IGS, MutS and LR DNA regions for the holotype and the redescribed colonies of *Pleurocorallium secundum*

B) The location of the PCR primers used in this study to amplify the IGS and MutS regions

C) The applied PCR protocols

**S2** Supporting results

DNA sequence alignments and NCBI GenBank accession numbers. Excel spreadsheet.

