## Supplementary material for "Trace-DNA from a century-old holotype specimen resolves taxonomic uncertainties: the case of the Hawaiian pink precious coral (*Pleurocorallium secundum*), a CITES-listed species used in jewelry": Lendvay_et_al_S1_Supporting_methods

for the article titled

by the authors

Bertalan Lendvay, Nadja V. Morf, Laurent L. Cartier, Michael S. Krzemnicki, Masanori Nonaka

**S1** Supporting Methods

A) PCR primers used for amplifying the mitochondrial IGS and MutS DNA regions for the holotype and the redescribed colonies of *Pleurocorallium secundum*

| Region | Primer name | Primer sequence | Amplicon length (bp, including primer sites) | Reference |
| --- | --- | --- | --- | --- |
| IGS | Pleuro-IGS1-F | 5'-ATC ACC ATA AAA CTA GCT CC-3' | 186 | this study |
| | Pleuro-IGS1-R | 5'-GGC GCA TTC AAA AGT CC-3' | | |
| | Pleuro-IGS2-F | 5'-CCT CTA CAA CAG TTG ATA GG-3' | 205 | this study |
| | Pleuro-IGS2-R | 5'-TCT CTA GTT AGG GTT TAC GC-3' | | |
| | Pleuro-IGS3-F | 5'-GAC TGG GCA GTT AGT GG-3' | 231 | this study |
| | Pleuro-IGS3-R | 5'-TAT GAT TAG TAG AAA ATA GCC AGC-3' | | |
| | Pleuro-MutS1-F | 5'-GAA CCA GAC TTA CTT TGG G-3' | 195 | this study |
| | Pleuro-MutS1-R | 5'-CCA TCT ATC TGT TAT TCA GC-3' | | |
| | *Coral*-ID-F | 5'-TACGYTCATAAATTATHCCT-3' | 189 | Lendvay et al. (2022) |
| | *Coral*-ID-R | 5'-AGATTTGCCATGGYACAGAA-3' | | |
| | Pleuro-MutS2-F | 5'-ATT GCT AGC CCT GAA TTT AC-3' | 140 | this study |
| | Pleuro-MutS2-R | 5'-GAC GAT TTT CAT AAG GGC CAA-3' | | |
| LR | LR-F | 5'-AAG TAG TGT CAC TCC CAA ACA-3' | 170 | Lendvay et al. (2020) |
| | LR-R | 5'-TGC AAA GAA GGA GAA CAA AAG G-3' | | |

B) The location of the PCR primers used in this study to amplify the IGS and MutS regions as mapped on the *Pleurocorallium konojoi* mitochondrial reference genome (NC_015406, (Uda *et al.* 2011)). Graphical output from Geneious Prime 2022.0.2 software

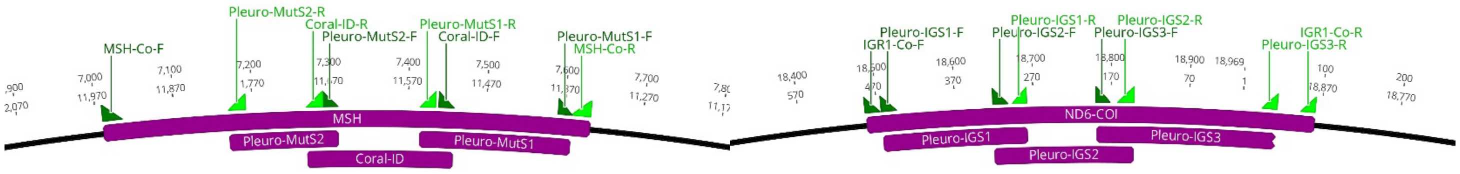


C) The applied PCR protocols

PCR reactions were performed in 25 µl final volume containing 1×Qiagen Multiplex PCR Master Mix (Qiagen) and 0.4 µM of the forward and reverse primers, respectively and 5 µl template DNA Thermal cycling commenced with enzyme activation at 95 ºC for 10 min followed by 50 cycles of denaturation 94 ºC for 45 sec, 53 ºC annealing for 1 min 30 sec and elongation at 72 ºC for 45 sec, and a final elongation of 72 ºC at 10 min was applied. PCR amplicons were purified with the AmPure XP bead system (Beckman Coulter) and quantified with Qubit fluorimeter device (Invitrogen).
